# Personalized characterization of diseases using sample-specific networks

**DOI:** 10.1101/042838

**Authors:** Xiaoping Liu, Yuetong Wang, Hongbin Ji, Kazuyuki Aihara, Luonan Chen

**Author notes:** These authors contributed equally to this work (Xiaoping Liu, Yuetong Wang).

## Abstract

A complex disease generally results not from malfunction of individual molecules but from dysfunction of the relevant system or network, which dynamically changes with time and conditions. Thus, estimating a condition-specific network from a sample is crucial to elucidating the molecular mechanisms of complex diseases at the system level. However, there is currently no effective way to construct such an individual-specific network by expression profiling of a single sample because of the requirement of multiple samples for computing correlations. We developed here with a statistical method, i.e., a sample-specific network method, which allows us to construct individual-specific networks based on molecular expression of a single sample. Using this method, we can characterize various human diseases at a network level. In particular, such sample-specific networks can lead to the identification of individual-specific disease modules as well as driver genes, even without gene sequencing information. Extensive analysis by using the Cancer Genome Atlas data not only demonstrated the effectiveness of the method, but also found new individual-specific driver genes and network patterns for various cancers. Biological experiments on drug resistance further validated one important advantage of our method over the traditional methods, i.e., we even identified those drug resistance genes that actually have no clearly differential expression between samples with and without the resistance, due to the additional network information.

## INTRODUCTION

One key to achieving personalized medicine is to elucidate molecular mechanisms of individual-specific diseases, which generally result from dysfunction of individual-specific networks/systems rather than malfunction of single molecules (Barabasi et al., 2011; Chen et al., 2009; Chen et al., 2010; Hood and Flores, 2012). In fact, it has been recognized that the phenotypic change of a living organism can seldom be fully understood by merely analyzing single molecules, and it is the relevant system or specific network that is ultimately responsible for such a phenomenon (Chen et al., 2009; Chen et al., 2010). With rapid advances in high-throughput technologies, applying molecular networks to the analysis of human diseases is attracting increasingly wide attention (Barabasi et al., 2011). A molecular network, e.g., a gene regulatory network, or a co-expression network, can be generally estimated by correlation coefficients of molecule pairs from expression or sequence data of multiple samples. Based on biological and clinical data, a number of network-based methods were proposed not only to identify disease modules and pathways but also to elucidate molecular mechanisms of disease development at the network level (Fang et al., 2012; Ideker and Krogan, 2012; Liu et al., 2012). To determine a person’s state of health, many studies have shown that network-based biomarkers, e.g., subnetwork markers (Ideker and Krogan, 2012; Liu et al., 2012), network biomarkers (Liu et al., 2014), and edge biomarkers (Zhang et al., 2014), are superior to traditional single-molecule biomarkers for accurately characterizing disease states due to their additional information on interactions and networks. In particular, an individual-specific network is considered to be reliable for accurately characterizing the specific disease state of an individual. It can be directly used to identify the biomarkers and disordered pathways, and further elucidate the molecular mechanisms of a disease for individual patients. However, it is generally difficult to obtain individual-specific networks (i.e., networks on an individual basis) because constructing an individual-specific network from expression data by traditional approaches requires multiple samples so as to evaluate correlations or other quantitative measures (Bhardwaj and Lu, 2005; De Bodt et al., 2009; Liu et al., 2012; Zhang et al., 2013) between molecules for each individual, which are usually not available in clinical practice and thus seriously limit their application in personalized medicine. In other words, although we can now obtain information of individual-specific differentially expressed genes or somatic mutations from expression or sequence data (Liu et al., 2013; Mendelsohn, 2013; Meric-Bernstam et al., 2013) of a single sample, there is still no effective methodology to construct the individual-specific network from such data of the single sample, which is the key personalized feature of each individual at a system level.

In this study, we developed a statistical method to construct an individual-specific network solely based on expression data of a single sample, i.e., a single-sample network or sample-specific network (SSN), rather than the aggregated network for a group of samples, based on statistical perturbation analysis of a single sample against a group of given control samples. In particular, we derived the SSN method to quantify the individual-specific network of each sample in terms of statistical significance in an accurate manner, which is the theoretical foundation of this method. Analyses of the Cancer Genome Atlas (TCGA) data with nine different cancers not only validated the effectiveness of our method, but also led to the following discoveries: (1) we found that there are several common network patterns in the same types of cancer, which, however, are not shared by other types of cancer; (2) personalized features of various cancers were characterized by SSNs, which in turn also revealed important regulatory patterns of driver genes in cancer; (3) individual somatic mutations for a sample were strongly correlated with its SSN on a single-sample basis, which was also validated by pathway enrichment and functional analysis; and (4) in contrast to the mutational driver genes, the functional driver genes, which functionally affect the occurrence and development of cancer, can be predicted from the hub genes of an SSN for an individual sample. As further applications of TCGA to big data, SSNs were used to predict individual driver mutations for various cancers solely based on gene expression without DNA sequence information, classify cancer phenotypes, and identify cancer subtypes by network biomarkers for accurate diagnosis and prediction of diseases in individuals, which all agree well with the experimental data.

Moreover, knockdown experiments validated our prediction of drug-resistant genes in the lung cancer cell line PC9. In contrast to traditional methods that are based on the differential gene expression between the samples with and without drug resistance, we further identified those drug-resistant genes that actually have no differential expression and thus are generally missed by traditional methods. Note that the SSN also means an individual-specific network in this study.

## RESULTS

### Constructing an SSN from a single sample

The SSN for each sample or individual is constructed based on statistical perturbation analysis of this sample against a group of given control samples. For this, we need to have expression data for a group of samples, which serve as the control or reference samples. As shown in Figure 1, by using this group of samples, we can construct the reference network by Pearson correlation coefficients (*PCCs*) (Figure 1A), i.e., compute the *PCC* of each pair of molecules as an edge with or without a template or background network. Generally, the reference network would have the common attributes of these reference samples. We then add the single test sample *d* to this group and construct another network by *PCCs*; this new network is called the perturbed network (Figure 1B). Thus, we can obtain the differential network between the reference and perturbed networks, which can clearly characterize the specific features of the additional sample *d* against this group. This differential network is referred to as the SSN of this new sample (Figure 1C). The key is how to quantify the statistical significance of each differential edge (i.e., differential *PCC)* in the differential network. Based on the analysis of perturbation statistics, we derived SSN theory (see next section) to accurately quantify each differential edge in the network for a single sample in terms of statistical significance, which is the theoretical foundation for this method (Figure 1C). All of the edges with significant differential correlations were used to constitute the SSN for the single sample *d* (Figure 1B and 1C). In this study, the functional association network with high confidence (confidence score ≥ 0.9) was used as the template or background network from the STRING database (http://string-db.org) that includes physical interactions, regulatory interactions, and the co-expression network of molecules, and all edges in the template network were measured by the *PCC* (Supplemental Information Note S2), which was calculated by the “SciPy” extension module (http://www.scipy.org/) of the Python programming language.

**Figure 1.**
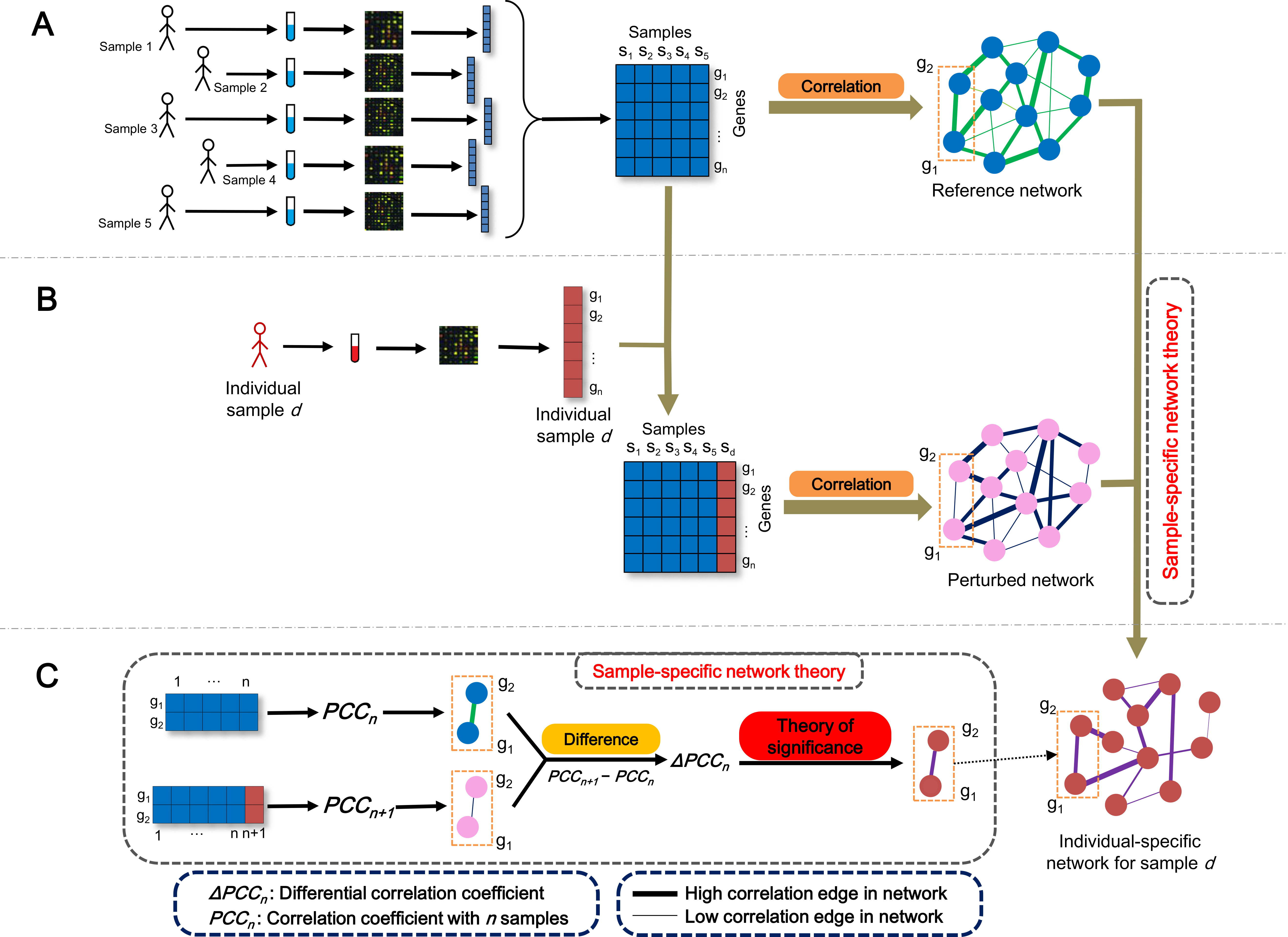
Flowchart for constructing an individual-specific network. (A) For a group of reference samples (*n* samples), a reference network can be constructed by the correlations between molecules based on expression data of this group of samples (multisamples), i.e., computing the *PCC*_*n*_ (the Pearson correlation coefficient of an edge in the reference network with *n* samples) of each pair of molecules as an edge in the network. Generally, the reference network has the common attributes of these reference samples. (B) A new sample d is added to the group, and the perturbed network with this additional sample is constructed in the same way by the correlation *PCC*_*n*+1_ of the combined data. The difference between the reference and perturbed networks is due to sample *d*. (C) The differential network is constructed by the difference of the corresponding edge between the reference and perturbed networks in terms of PCC, i.e., Δ*PCC_n_* = *PCC_n+1_* − *PCC_n_* for each edge. Based on sample-specific network theory, we can quantify the significance of each edge, i.e., Δ*PCC_n_* in the network. The SSN for sample d is constituted by those edges with significant Δ*PCC_n_*.

### Theoretical foundation of the SSN

For the differential network, each edge is a differential *PCC* (Δ*PCC)*, and we provide a quantitative measure to evaluate its statistical significance. Assuming that there are *n* samples for the group of the given reference samples, we refer to the *PCCs* of an edge in the reference network with *n* samples and the perturbed network with *n + 1* samples (due to one additional test sample) as *PCC_n_* and *PCC_n+1_*, respectively. Then, the Δ*PCC* of the edge between reference and perturbed networks is Δ*PCC* = *PCC_n+1_* — *PCC_n_*. We can show that the mean and standard deviation of Δ*PCC_n_* for the population are *µ*_Δ*PCC*_=*O*(1/*n*^2^) ≈ 0 and 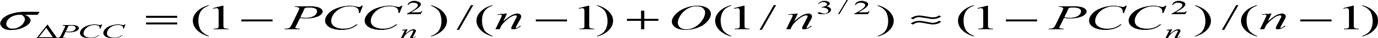 with a large *n*, where *O(1/n)* implies the term with the order of *1/n* (Supplemental Information Note S2). It is well known that the *p* value of *PCC_n_* can be evaluated by Student’s *t* test with *n* — 2 degrees of freedom (Supplemental Information Note S3), i.e.,

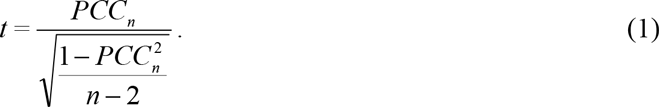

In this work, we can theoretically further show that the *p* value of such Δ*PCC* follows a new type of symmetrical distribution defined as volcano distribution in this paper (Supplemental Information Note S4), whose tail regions are similar to those of the normal distribution. Hence, the statistical hypothesis test *Z*-test (or *U*-test) can be used to evaluate the significance level of each Δ*PCC* because of the central limit theorem (Rice, 2007). The null hypothesis is that the Δ*PCC_n_* is equal to the population mean of Δ*PCC_n_,* and thus we have

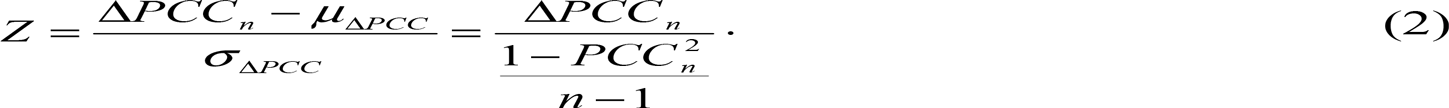

From Eqn. (2), the *p* value for the edge can be obtained from the statistical *Z* value (Figure 2A). If the *p* value of *Z*-test is less than 0.05, then the Δ*PCC* or this edge is significant and there is such an edge in the perturbed or differential network. From Eqn. (2), we can see that the *p* value of Δ*PCC_n_* depends on the number of samples in the reference group and also the correlation of the reference edge as well as Δ*PCC_n_*, and thus different numbers of control samples and the correlation of the edge will yield a different significance of the edge in the SSN, even with the same value of the differential *PCC* (Δ*PCC_n_*). Note that the *p* value of *PCC* depends on the *t* distribution with only two parameters, *n* and *PCC_n_* (see Supplementary Information).

**Figure 2.**
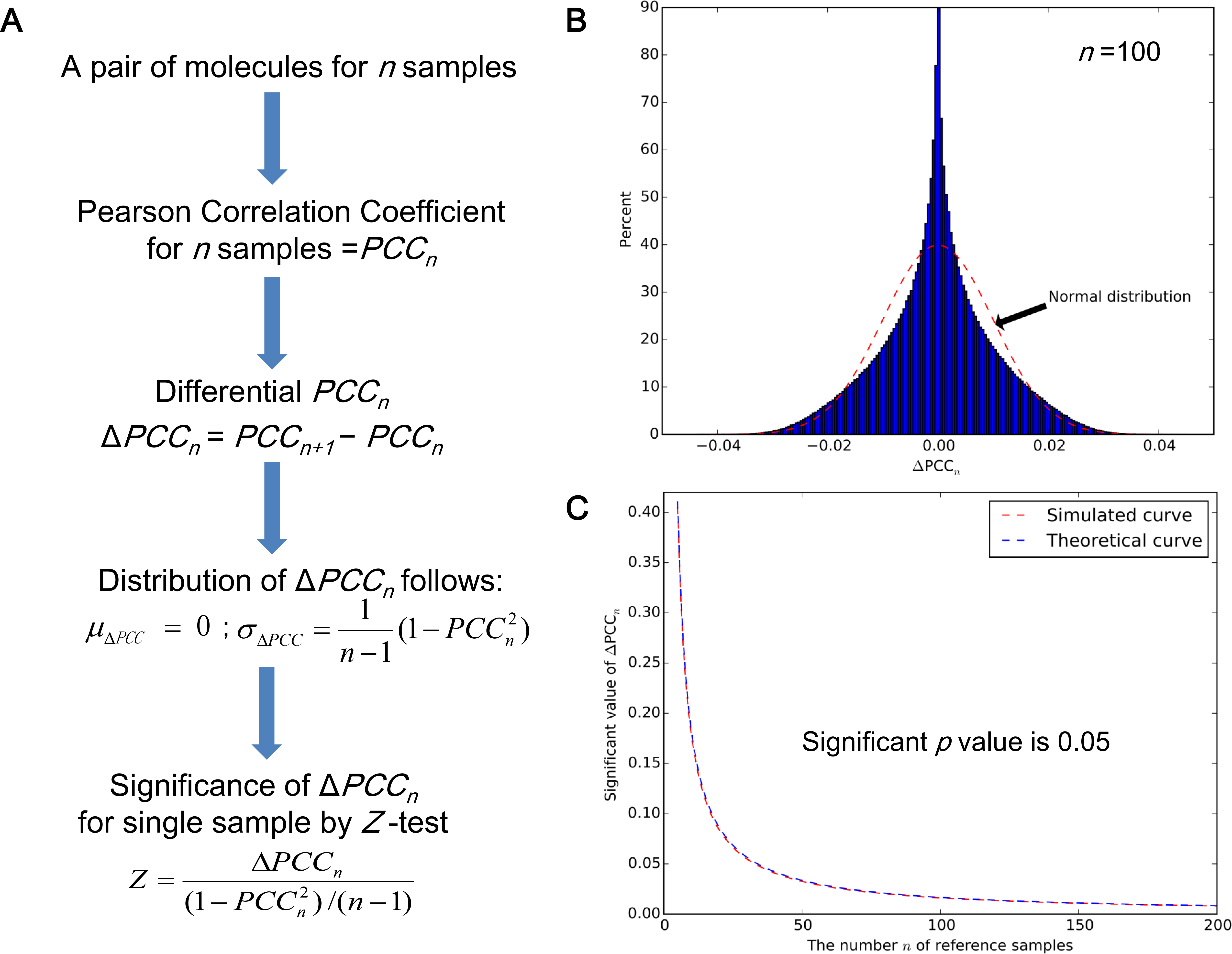
The significance of a differential Pearson correlation coefficient (Δ*PCC*) or an edge. (A) The theoretical result to evaluate the significance of PCCn by Eqn. (2), (B) the distribution of Δ*PCC_n_* numerically obtained by random simulation (*n* = 100), (C) the significant value of Δ*PCC_n_* evaluated by the numerical simulation (i.e., from the distribution of the random simulation) and the theoretical result (i.e., from Eqn. (2)). Δ*PCC_n_* in the area above the curve is statistically significant with a *p* value of < 0.05. Clearly, the simulated curve (red color) and theoretical curve (blue color), i.e., the values of PCCn for the random simulation and the theoretical calculation of Eqn. (2) with a *p* value of 0.05 are almost identical with little difference, which well validates Eqn. (2) of the sample-specific network method.

For validation of Eqn. (2), we randomly generated two series of reference numbers (i.e., the expression of two molecules) to estimate their correlation as an edge based on multivariate normal distribution with the different correlation *PCC_n_* = 0, 0.1 to 0.9 of the two series of numbers by the “Numpy” package (http://www.numpy.org/). The number or length *n* of the two series was changed from 5 to 200 (i.e., the number of the reference samples). For every pair of *n* and *PCC_n_*, the random digital simulation was repeated 2,000,000 times, where the value of Δ*PCC_n_* with a *p* value of 0.05 in the two-tails area was selected from every distribution of simulation, i.e., the significant value (Figure 2C, red line, and Figure S2). As shown in Figure 2B (*n* = 100), the distribution of Δ*PCC_n_* follows a new type of distribution defined as volcano distribution, whose tail areas are similar to those of a normal distribution in a random condition. At the same time, the significant value of Δ*PCC_n_* with the *p* value of 0.05 can also be obtained from the theoretical calculation (Figure 2C, blue line, and Figure S3), i.e., Eqn. (2), where Δ*PCC_n_* in the area above the curve is statistically significant with a *p* value of < 0.05 (Figure 2C). The simulated and theoretical curves, i.e., the values of Δ*PCC_n_* for the random simulation and the theoretical calculation of Eqn. (2) with the *p* value of 0.05 are almost identical (Figure 2C, and in particular, Figures S1 and S2 for all *n* and *PCCn)* with little difference, which well validates Eqn. (2) of the SSN method.

### SSNs reveal common network patterns for cancers at molecular level

We chose the datasets of nine different types of cancers from TCGA (http://cancergenome.nih.gov/) database (Table S1 and Supplemental Information Note S1) with gene expression profiling and matched clinic information. For each type of cancer, 8–17 normal samples were selected as the reference samples (Table S1), and every single cancer sample was used to construct its SSN (Figure 1 and Table S2). We collected the expression and sequence data of the nine cancers, and detailed information for the numbers of control and cancer samples is given in Table S1. The SSN for each sample was constructed for all cancer datasets, and only the correlation-lost edges (see Note S10), whose correlations in absolute values decrease from the reference to this sample, were chosen to perform the following analysis. We focused here on the analysis of the sample-specific subnetworks with correlation-lost dges related to tumor protein p53 (TP53), which is a crucial gene in cancer, to demonstrate the power of this analysis for characterizing the personalized features (Figure 3). The subnetwork of TP53 is composed of genes directly connected with TP53, or its first-order neighboring genes. Figure 3A shows the subnetworks of TP53 from four breast invasive carcinoma (BRCA) samples, which clearly characterize their individual features at the network level. Although most edges in the four subnetworks of TP53 are different from each other, the associations between TP53 and both MUC1 and CCNB1 exist in all four subnetworks (Figure 3A). In fact, by analyzing subnetworks of TP53 for all 761 samples in breast cancer, we found that there is an association between TP53 and MUC1 in 65.18% of breast cancer samples, and an association between TP53 and CCNB1 in 64.52% of breast cancer samples File S1). In other words, for 65.18% of individuals with breast cancer, the correlation between TP53 and MUC1 has a significant loss in the cancer status, and the correlation between TP53 and CCNB1 has been lost in the cancer status for 64.52% of individuals with breast cancer. MUC1 is a known marker for breast cancer (Wreschner et al., 1994), and it associates with TP53 (Wei et al., 2005, 2007) and plays an important role in breast cancer (Gimmi et al., 1996; Kufe, 2013). CCNB1 is an important known marker for breast cancer and has important implications for cancer prognosis (Ding et al., 2014; Suzuki et al., 2007).

**Figure 3.**
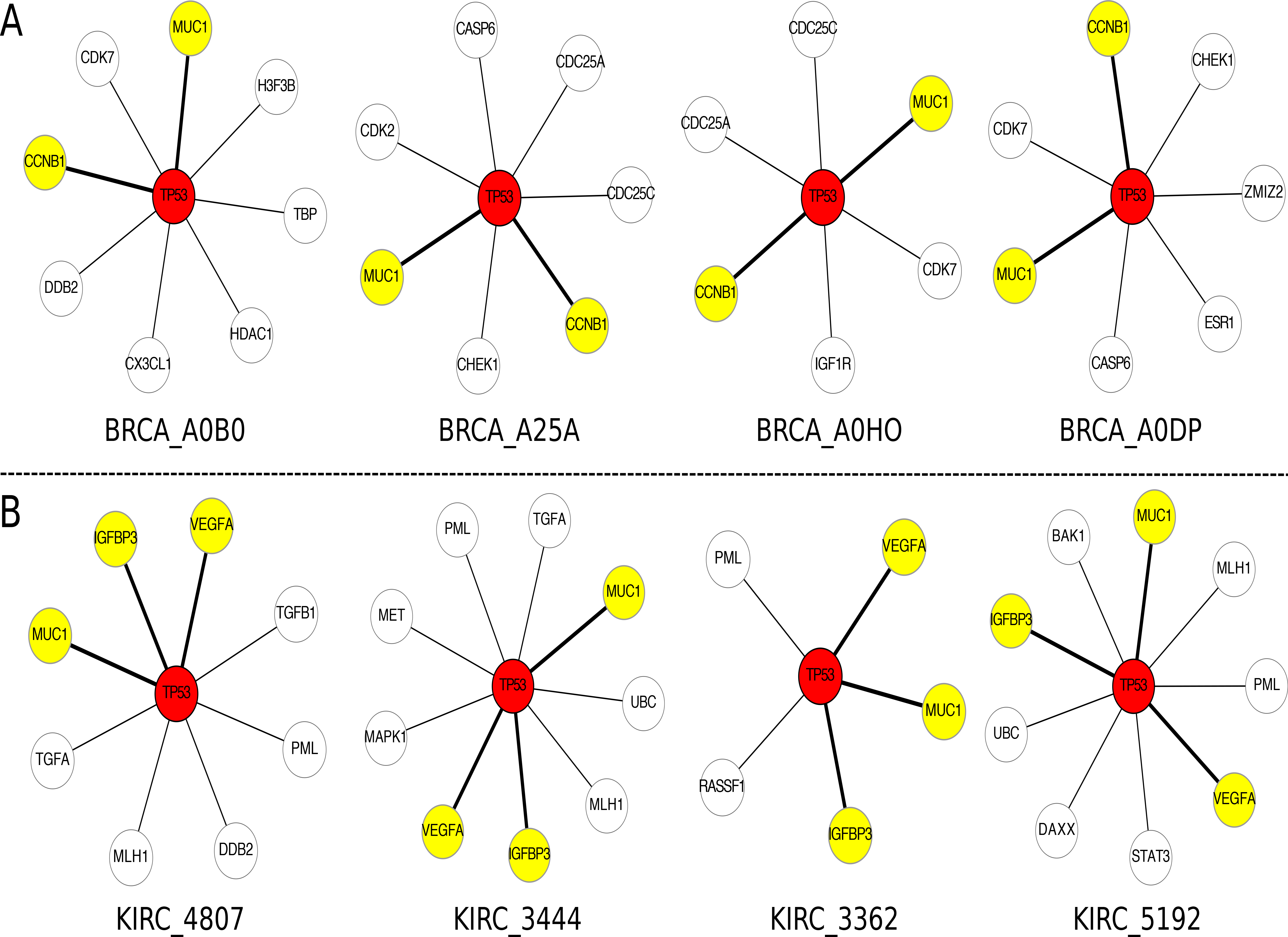
Individual-specific networks characterize personalized features and also reveal common network patterns for cancers. (A) The four individual-specific subnetworks of tumor protein p53 (TP53) from four samples for breast invasive carcinoma (BRCA). The numbers of the connections with TP53 for the four samples are respectively 8, 7, 6, and 7, and the genes linked to TP53 are also different in the four samples, i.e. DDB2 for BRCA_A0B0, CHEK1 for BRCA_A25A, IGF1R for BRCA_A0HO, and ESR1 for BRCA_A0DP are unique genes for the respective subnetworks of the four samples. However, MUC1 and CCNB1 (yellow color) are common genes appearing in the four subnetworks. Actually, we found that 65.18% of BRCA samples include a significant connection (bold lines) between TP53 and MUC1, and 64.52% of BRCA samples include a significant connection (bold lines) between TP53. i.e., these connections have a significant loss of correlation between BRCA and normal samples, and is the common network pattern related to TP53. (B) The individual-specific subnetworks of TP53 from four samples for kidney renal clear cell carcinoma (KIRC). There are different targets and numbers connected with TP53 in the four samples of KIRC. Three genes, VEGFA, IGFBP3, and MUC1 (yellow color) appeared in all four samples (bold lines). Actually, we found that 94.74% of KIRC samples have a significant loss of correlation for connection between VEGFA and TP53, 92.34% of KIRC samples have a significant loss of correlation for connection between IGFBP3 and TP53, and 87.32% of KIRC samples have a significant loss of correlation for connection between MUC1 and TP53, which are the common network patterns related to TP53.

By analyzing the gene expression of the four breast cancer samples, we found that TP53 is significantly upregulated only in two (BRCA_A0B0 and BRCA_A25A) of the four samples, as compared with its expression in the reference samples, and the gene expression of TP53 was not significantly changed in the other two samples (BRCA_A0HO and BCRA_A0DP) compared with the reference samples, but MUC1 was significantly upregulated in all four samples. The correlation between TP53 and MUC1 is a positive value relative to the reference samples, and the correlation coefficient is significantly decreased in the four samples. Thus, we called the relationship between TP53 and MUC1 in breast cancer +−UU (Positive Correlation Decreases due to X Upregulation and Y Upregulation, TP53 as Y, see Class–5 in Note S11 and Table S6) for sample BRCA_A0B0 and BRCA_A25A, and +−^*^N (Positive Correlation Decreases due to X Upregulation and Y No change, TP53 as Y, see Table S6) for sample BRCA_A0HO and BRCA_A0DP (Supplemental Information Notes 11). The gene CCNB1 is also significantly upregulated in all four samples. The correlation between TP53 and CCNB1 is a positive value based on the reference samples, and the correlation coefficient is significantly decreased in the four samples. Thus, we called the relationship between TP53 and CCNB1 +−UU for samples BRCA_A0B0 and BRCA_A25A, and +−^*^N for samples BRCA_A0HO and BRCA_A0DP (TP53 as Y, Supplemental Information Note 11 and Table S6).

On the other hand, in the subnetworks of TP53 for kidney renal clear cell carcinoma (KIRC), the network patterns or targets of TP53 are also diverse and individual dependent (Figure 3B), but there are three consistent edges connected with VEGFA, IGFBP3, and MUC1 in all four subnetworks of TP53. By analyzing all individual-specific subnetworks of TP53 for 418 kidney cancer samples, we found the edge with VEGFA in 94.74% of SSNs, the edge with IGFBP3 in 92.34% of SSNs, and the edge with MUC1 in 87.32% of SSNs (File S1). This means that the edge between TP53 and VEGFA among 94.74% of samples, the edge between TP53 and IGFBP3 among 92.34% of samples, and the edge between TP53 and MUC1 among 87.32% of samples have significant loss of correlation, or these associations in most kidney cancer samples suffer from significant loss compared with normal samples, i.e., they are the common network patterns related to TP53 in kidney cancer. The VEGFA gene is an important growth factor acting in kidney cancer (Luan et al., 2003; Rennel et al., 2007), the IGFBP3 gene is a cell growth factor and an important marker in kidney cancer (Cheung et al., 2004; Chuang et al., 2008), and the MUC1 gene affects invasive and migratory properties of kidney cancer cells and is a potential therapeutic target (Aubert et al., 2009). From the above analysis, the abnormal interactions of TP53 with VEGFA, IGFBP3, and MUC1 are considered to be potential factors contributing to kidney cancer.

By analyzing the gene expression of the four kidney cancer samples, we found that in three (KIRC_2444, KIRC_3362 and KIRC_5192) of the four samples TP53 is significantly upregulated relative to the reference samples, and the gene expression of TP53 did not significantly change only in sample KIRC_4807, and the VEGFA gene is significantly upregulated in all four samples. The correlation between TP53 and VEGFA is a negative value based on the reference samples, and the correlation coefficient is increased (loss of a negative correlation) in the four samples. Thus, we called the relationship between TP53 and VEGFA −+UU (Negative Correlation Increases due to X Upregulation and Y Upregulation, TP53 as Y) for samples KIRC_2444, KIRC_3362 and KIRC_5192, and −+^*^N (Negative Correlation Increases due to X Upregulation and Y No change, TP53 as Y) for sample KIRC_4807. The IGFBP3 gene is significantly upregulated in all four samples, the correlation between TP53 and IGFBP3 is a positive value based on the reference samples, and the correlation coefficient is decreased (loss of a positive correlation) in all four samples. Thus, we called the relationship between TP53 and IGFBP3 +-UU for samples KIRC_2444, KIRC_3362 and KIRC_5192, and +−^*^N for sample KIRC_4807. The gene MUC1 is downregulated in all four samples, the correlation between TP53 and MUC1 is a positive value based on the reference samples, and the correlation coefficient is decreased (loss of a positive correlation) in all four samples. Thus, we called the relationship between TP53 and MUC1 in kidney cancer +-DU (Positive Correlation Decreases due to X Downregulation and Y Upregulation, TP53 as Y) for samples KIRC_2444, KIRC_3362 and KIRC_5192, and +−^*^N (Positive Correlation Decreases due to X Downregulation and Y No change, TP53 as Y) for sample KIRC_4807 (Supplemental Information Notes 10 and 11 and Table S6).

Here, the loss of correlation means that the correlation in terms of the absolute value decreases from the reference samples.

### SSNs characterize personalized features and also reveal different regulatory patterns of driver genes in cancer

Each cancer sample has its own individual-specific pathogenesis. We used the individual-specific subnetworks related to the driver mutation gene (DMG) TP53 as an example to show the personalized features and regulatory patterns for various cancers.

The samples were selected from stomach adenocarcinoma (STAD), BRCA, and glioblastoma multiforme (GBM), and the subnetworks of TP53 are shown in Figures 4 and S3–S5, where each network is the individual-specific subnetwork for one sample. We classified the edges into two types, i.e., one is the correlation-gained edges (red lines) whose *PCCs* are increased from the reference network with positive correlation coefficients or decreased from the reference network with negative correlation coefficients, and another is the correlation-lost edges (black lines) whose *PCC*s are decreased from the reference network with positive correlation coefficients or increased from the reference network with negative correlation coefficients. For STAD, there are almost equal numbers of the correlation-gained and correlation-lost edges in the TP53 subnetwork (one sample in Figure 4A and four samples in Figure S4). Actually, on average from all 183 STAD samples, there are 28.37 increasing edges and 28.89 decreasing edges, which means that TP53 as a driver gene affects both the correlation-gained and correlation-lost edges in STAD. For BRCA, the correlation-lost edges are obviously greater than the correlation-gained edges in the TP53 subnetwork (one sample in Figure 4B and four samples in Figure S5). On average from all 761 BRCA samples, there are 2.22 correlation-gained edges and 26.34 correlation-lost edges; this implies that the interaction partners of TP53 are significantly reduced in BRAC, which is a general feature of BRAC. In contrast, for GBM, the correlation-gained edges are much greater than the correlation-lost edges for TP53 subnetworks (one sample in Figure 4C and four samples in Figure S6). On average from 333 GBM samples, there are 120.24 correlation-gained edges and 24.02 correlation-lost edges for the driver gene TP53, which characterizes the regulatory patterns of GBM, and means that interaction partners of TP53 are significantly increased. TP53 may promote tumor onset by mainly inducing the correlation-gained edges in GBM.

**Figure 4.**
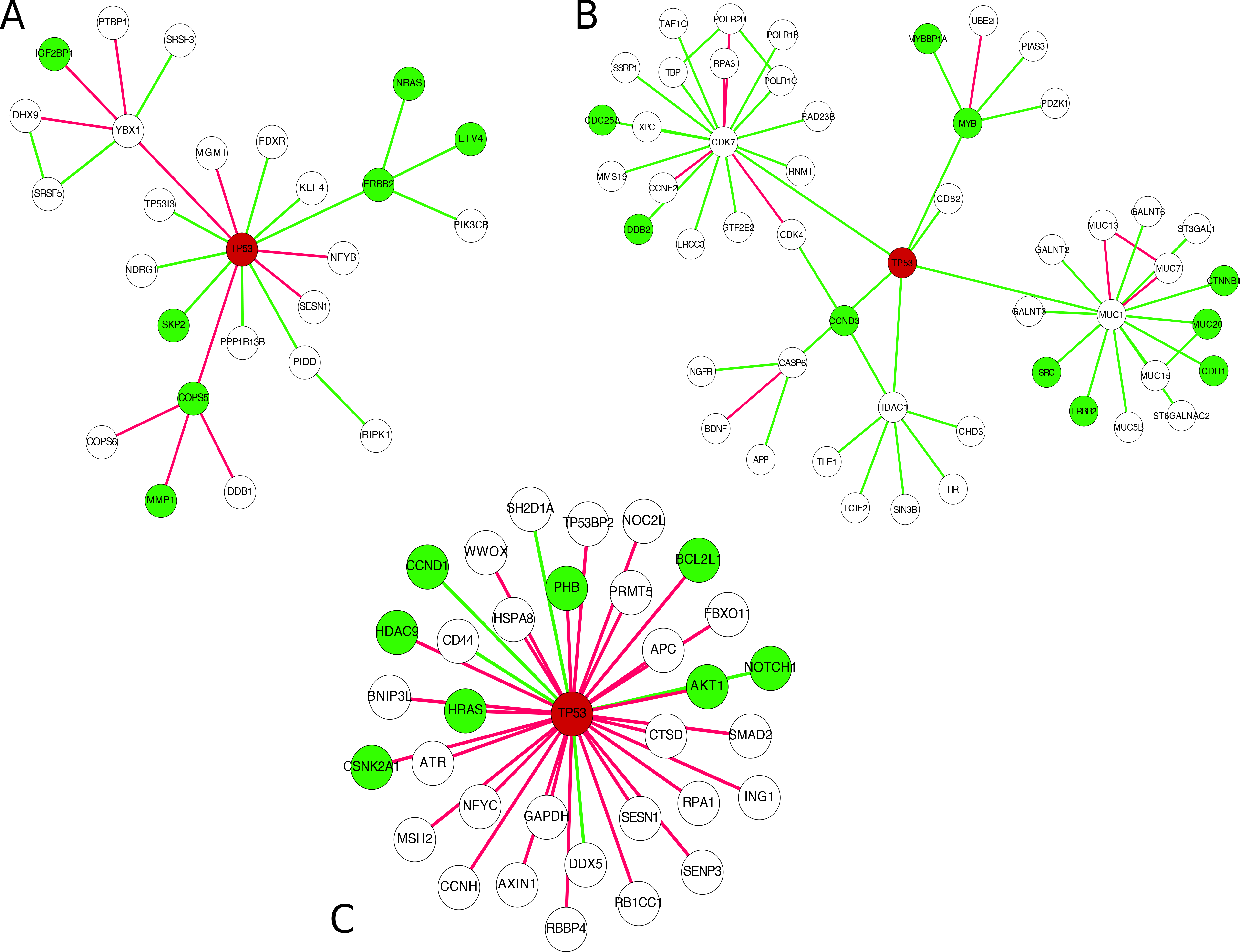
Individual-specific networks related to driver gene tumor protein p53 (TP53) reveal the regulatory patterns in different cancers. (A) The individual-specific subnetwork of TP53 in sample 4255 from stomach adenocarcinoma. (B) The individual-specific subnetwork of TP53 in sample A0CV from breast invasive carcinoma. (C) The individual-specific subnetwork of TP53 in 2574 from glioblastoma multiforme. The network analyses related to the driver gene TP53 not only reveal the personalized features of each sample but also indicate that each cancer type has a specific regulatory pattern. Red and green lines represent the correlation-gained and correlation-lost edges, respectively. A green node indicates an oncogene from NCBI (http://www.ncbi.nlm.nih.gov/).

### SSNs validated by literature

In addition to the descriptions in the preceding section, our findings for many other differential associations or regulations between normal and cancer samples are also consistent with previous reports. ESR1 (estrogen receptor 1) is an important factor for the pathological process of breast cancer. We found a significant loss of correlation between TP53 and ESR1 in 42.97% of breast cancer samples compared with the normal reference samples. Indeed, the ESR1 gene was found to be upregulated by TP53 in the breast cancer cell line MCF–7 (Angeloni et al., 2004; Rasti et al., 2012). The overexpression of ESR1 could cause the observed loss of correlation in breast cancer samples.

TP63 (tumor protein p63) is a member of the p53 family of transcription factors, and the correlation between TP63 and TP53 is usually low in normal lungs. A recent study showed a high frequency of co-expression between TP53 and TP63 in lung squamous cell carcinoma, but not in lung adenocarcinoma (LUAD) (Choy et al., 2013). Coincidentally, our analysis showed that despite low or even no correlation between TP53 and TP63 in the reference and normal samples, 74.2% of lung squamous cell carcinoma samples showed a significant gain in correlation compared with the reference samples, whereas only 6.5% of LUAD samples showed a significant correlation variation compared with the reference samples.

The CDC25C (cell division cycle 25C) gene is involved in the regulation of cell division. It is well known that CDC25C regulates cellular entry into mitosis in the G2/M phase (Moon et al., 2010), and its expression is also regulated by TP53 (St Clair et al., 2004). Hence, the abnormal regulation between TP53 and CDC25C may disrupt the regulation of CDC25C by TP53, and render the cell cycle and cell division out of control. The correlation between TP53 and CDC25C is 0.52 (the *p* value of significant *PCC* is 0.04) in the reference samples, which is significantly high. However, 52.04% of breast cancer samples showed a significant loss of correlation (correlation coefficient decreasing) between these two genes compared with the reference samples. This result is consistent with those of previous studies, i.e., it was found that CDC25C expression is regulated by TP53 (St Clair et al., 2004) in normal samples, but in breast cancer its expression and that of its splice variants are not dependent on TP53 (Albert et al., 2012).

### SSNs validated as personalized features by disease gene enrichment, individual somatic mutations, functional analysis, and pathway enrichment

Somatic mutation genes (SMGs) of cancer that provide individual-specific information for each sample (Blau and Liakopoulou, 2013) can be used to validate SSNs and our method (Blau and Liakopoulou, 2013). Specifically, we measured the relationship between the SSN and the SMGs (Supplemental Information Note S5) in the same sample to validate the sample specificity of each SSN. The topological distance (Methods and Supplemental Information Note S6) and functional distance (Methods and Supplemental Information Note S7) between an SSN and a set of SMGs were used to measure their relationships for every sample in various cancer datasets. As shown in Figure 5A, the topological distance between the SSN and corresponding SMGs is significantly small in more than 99% of samples on average in each cancer, i.e., they are significantly related to each other, which implies that the SSN is indeed sample specific and reflects the personalized features. From a functional viewpoint, there are more than 72% of samples on average in all cancers in which each SSN and the corresponding somatic mutations are significantly related in terms of the functional distance of gene ontology (GO) annotations (Figure 5B and Table S3), and more than 63% of samples on average of all cancers, in which each SSN and the corresponding somatic mutations are significantly related in terms of the functional distance in Kyoto Encyclopedia of Genes and Genomes (KEGG) pathways (Figure 5B and Table S3). In most of the samples, the SSN and somatic mutations are also significantly related in terms of functional distance in the BBID (Biological Biochemical Image Database, http://bbid.grc.nia.nih.gov/) and BIOCARTA (http://www.biocarta.com/) pathways (Table S3). In addition to the somatic mutations, the driver mutations of cancer (Supplemental Information Note S5) are also individual-specific mutations, and currently, 125 DMGs have been determined for cancer (Vogelstein et al., 2013). Then, instead of using SMGs, we similarly performed another analysis using the overlapped genes between the 125 DMGs and SMGs as a set of the driver genes in each sample for measuring the relationships. Figures 5A–C show that the results for the driver genes are similar to those for the SMGs, but are more significant in the functional distance analysis (Figure 5C). In particular, comparing with 63% of samples on average as shown in Figure 5B, there are more than 94% of samples on average in which each SSN and the DMGs are significantly related in terms of KEGG pathways (Figure 5C and Table S3). These results indicate that the SSN is indeed sample specific and characterizes the personalized features of each sample at the network level.

**Figure 5.**
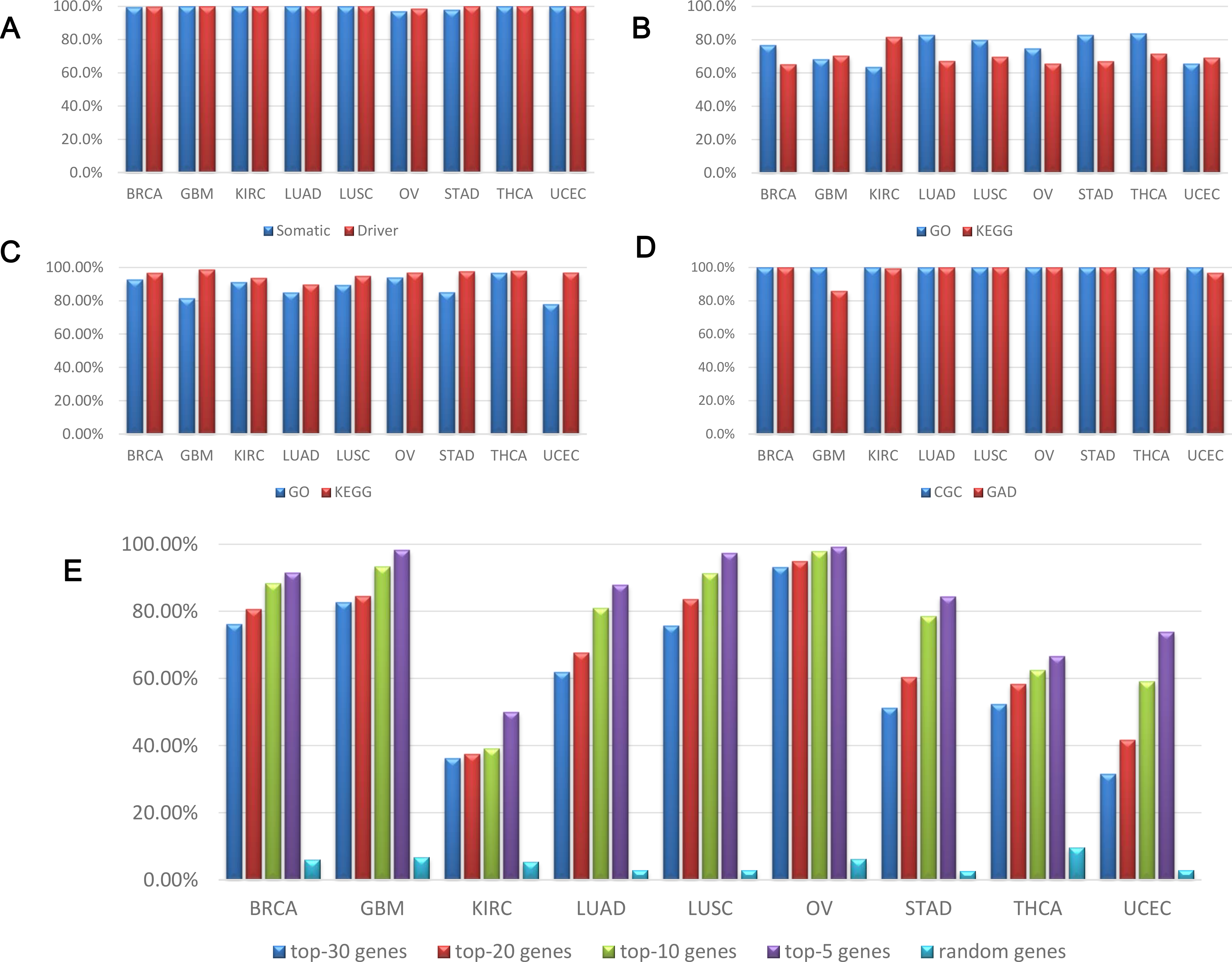
Validating individual-specific networks and predicting driver mutation genes in different cancers. (A) Topological distance between each sample-specific network and (somatic and driver) mutation genes in the same sample. (B) Functional distance between each SSN and somatic mutation genes in the same sample. (C) Functional distance between each SSN and driver mutation genes in the same sample. (D) Gene enrichment for each SSN in the Cancer Gene Census (CGC) and Genetic Association Database (GAD). The results in A-D show that each SSN is significantly related to the mutations of the same sample in terms of topological and functional distances, and indeed characterizes the personalized features of the individual. (E) Predicting individual driver mutation genes by each SSN in various cancers. The rate that a somatic mutation gene with a high degree is also a driver mutation gene increases in each SSN, and thus the accuracy of the prediction increases with the degree.

The known cancer genes can be downloaded from the CGC (Cancer Gene Census, http://cancer.sanger.ac.uk/cosmic/census) database and the GAD (Genetic Association Database, http://geneticassociationdb.nih.gov/), and the enrichment of an SSN relative to the known cancer genes can be used to validate its functional specificity (Supplemental Information Note S8). The result shows that the known cancer genes in the CGC database are significantly enriched in all of the SSNs in the cancer datasets (Figure 5D). For genes in GAD, we chose the cancer-associated genes, and then evaluated the significance of enrichment for each SSN (Figure 5D and Supplemental Information Note S8); our results clearly show that the corresponding cancer genes are significantly enriched in most SSNs.

### SSNs predict individual driver mutations for cancer solely based on gene expression without DNA sequence information

A hub node in an SSN is a gene that is highly connected with other genes (a gene with a high degree or with many links). Generally, the higher the gene degree in an SSN, the greater the variations or changes in the regulation related to this gene from normal to tumor samples, i.e., a gene with a high degree is a gene with large variations in interactions on a network level. Thus, a high-degree gene in the SSN is more likely to be a DMG for cancer. Based on this hypothesis, we predicted the DMGs for the genes with different degrees in SSNs (Supplemental Information Note S9), and found that the higher the gene degree, the more likely this gene is a DMG (Figure 5E). We conducted the computation for the top 30, 20, 10, and 5 highest degree genes for every SSN, and the rate that a gene with the high degree is also a DMG was calculated for every cancer by further checking the somatic mutation data of the genes. Clearly, from the top 30 highest degree genes the rate is monotonically increased (Figure 5E). For example, within the top 30 highest degree genes, about a half of those SMGs (51.23% rate) are DMGs for stomach cancer (STAD) in Figure 5E. With the top 5 highest degree genes, 84.44% of those SMGs are DMGs. However, in the random condition, only 2.71% of SMGs are DMGs in STAD (Figure 5E). Clearly, these results indicate that the hub genes in an SSN are strongly related to the driver mutations in the same sample, and thus can be used to predict the potential driver genes (including driver mutation and driver non-mutation genes, also called functional driver genes) on an individual basis for each sample, even without its DNA sequence information. As shown in Figure 5E, the accuracy of the prediction of the DMGs increases with the degree of the hubs in SSNs. These results also imply that high-degree genes in the network are high-risk genes that are more likely to be related to the tumor onset than other genes.

The potential driver genes for each sample (File S2) were obtained from the top 10 high-degree genes in the SSN of each sample. The functional enrichment analysis was performed for the potential driver genes of each sample using known DMGs and oncogenes from NCBI (http://www.ncbi.nlm.nih.gov), and we found that the known DMGs (Vogelstein et al., 2013) and oncogenes were significantly enriched (Figure S7). The significant enrichment of the known DMGs and oncogenes in the potential driver genes of most samples for every cancer clearly indicates that the SSN method is valid for predicting driver genes of cancer solely based on gene expression without DNA sequence information. In other words, the SSN method provides an effective strategy for personalized medicine, enabling the prediction of potential driver genes or potential oncogenes for a specific patient based on single-sample data.

### SSNs classify phenotypes of cancer and identify subtypes of cancer by network biomarkers for accurate diagnosis

Molecular networks are reliable forms to accurately characterize complex diseases, in contrast to individual molecules. Many network-based approaches have been proposed to extract a discriminative gene set as a biomarker for the classification of samples with distinct phenotypes by considering network information, but when diagnosing a new sample, such a gene set is simply used in a similar way to traditional molecular biomarkers without effectively exploiting the network information of that sample. Thus, those biomarkers are not network biomarkers but essentially molecule biomarkers. In contrast, our method can construct the SSN for each sample, and therefore opens a new way for diagnosing a single sample by network biomarkers, i.e., diagnose or classify each sample by the SSN/subnetwork/edges.

We first used hierarchical clustering to classify the normal and tumor samples, by using the top 5 differentially expressed genes (i.e., node biomarkers, by the traditional method) and the top 5 differential edges or Δ*PCC*s for SSNs (i.e., edge biomarkers, by our method) as the biomarkers. The edge biomarkers are clearly superior to node markers in terms of the accuracy of the classification of normal and tumor samples (STAD) by hierarchical clustering, i.e., there are only 4 samples for edge biomarkers but 29 samples for node markers, which were wrongly classified (top 5 edge biomarkers with accuracy of 98.1% in Figure 6A, and top 5 node biomarkers with accuracy of 86.5% in Figure 6B).

**Figure 6.**
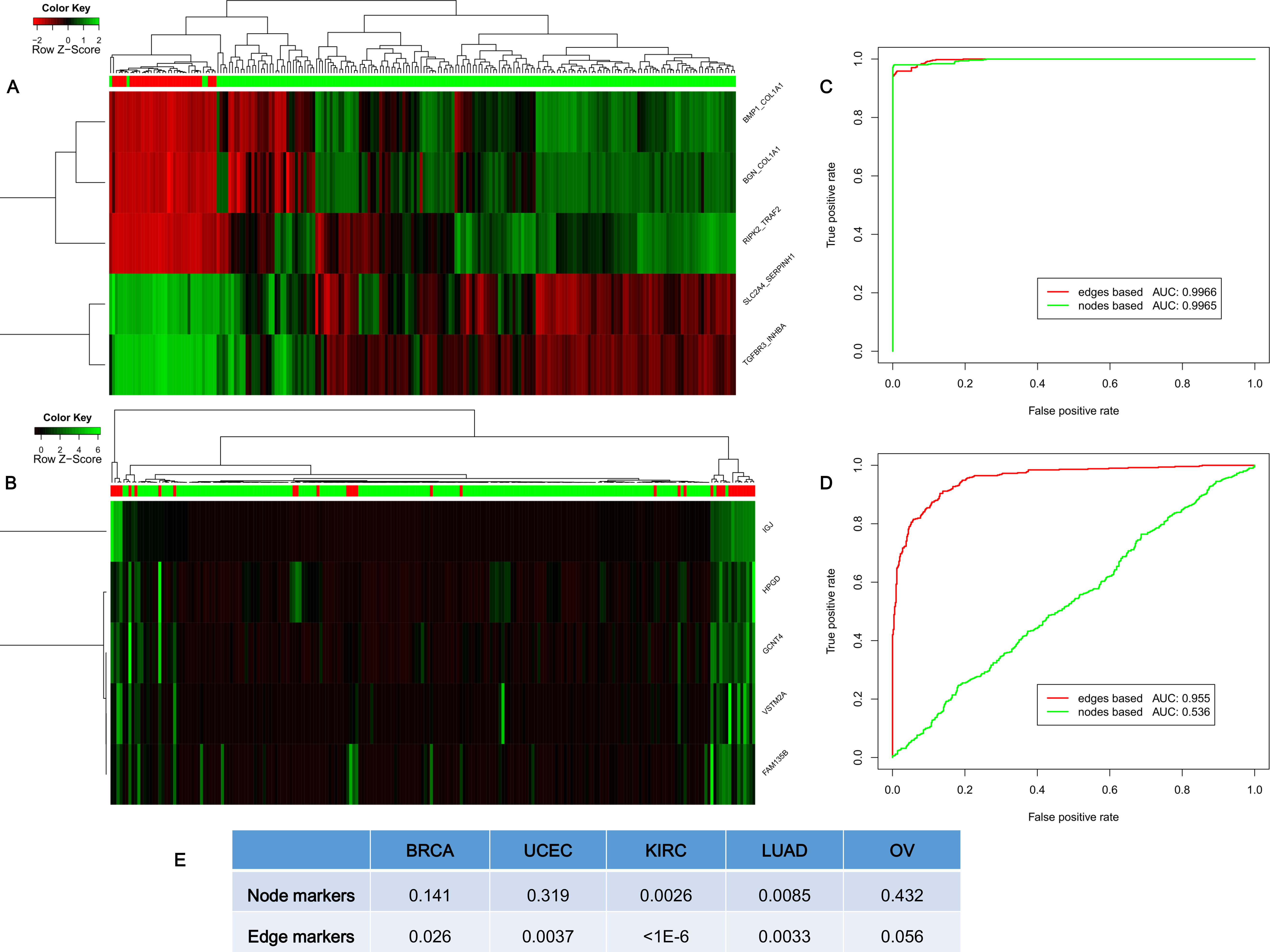
Classifying phenotypes and cancer subtypes. (A) The classification of cancer (183 samples, green bar) and normal (33 samples, red bar) samples by hierarchical clustering of the edge biomarkers (top 5 differential edges or differential Pearson correlation coefficients [Δ*PCCs*] by our method) for stomach adenocarcinoma (STAD) with 98.1% accuracy. (B) The classification of normal and cancer samples by hierarchical clustering of the node biomarkers (top 5 differential genes by the traditional method) for STAD with 86.5% accuracy. (C) The classification result of using the top 5 differential genes and edges for lung adenocarcinoma (LUAD). (D) The classification result of using the bottom 5 differential genes and edges for LUAD. (E) The log-rank *p* value of the survival curve for the subtyping in breast invasive carcinoma (BRCA), uterine corpus endometrial carcinoma (UCEC), kidney renal clear cell carcinoma (KIRC), LUAD and ovarian serous cystadenocarcinoma (OV). We used the top 100 variable genes as node biomarker for subtyping cancers (the traditional method), and top 100 variable edges (or Δ*PCCs*) as edge biomarkers for subtyping cancers (our method). All results show that individual-specific subnetworks or edge biomarkers are superior to the traditional node or molecular biomarkers in terms of classification and subtyping.

We then used the SVM (supporting vector machine) model with 5-fold cross-validation to classify the phenotypes or samples for various cancers. When we chose the top 5 differentially expressed genes and edges as the biomarkers to classify the normal and tumor samples, we found that the AUC for edge biomarkers and node biomarkers are respectively 99.66% and 99.65% for LUAD, and thus the accuracies of both are very similar and high (Figure 6C LUAD, and Figure S8 for the other eight cancers). However, for the same data of LUAD, when we chose the bottom 5 differential-expression genes and edges as the biomarkers to classify the two phenotypes, the AUC for edge biomarkers is still as high as 95.5% but the AUC for node biomarkers is significantly reduced to 53.6%, which implies that the edge biomarkers are robust and have synergetic power in the classification, compared with the node biomarkers (Figure 6D for LUAD, and Figure S8 for the other eight cancers).

Subtyping cancer is an important topic in recent cancer research, and most subtyping methods mine the information based on expression of genes, rather than individual networks. Here, although a sophisticated algorithm may considerably improve the accuracy, we used a simple subtyping method (Tothill et al., 2008) to identify the potential subtypes in different cancers separately based on gene expression data (i.e., node biomarkers, top 100 variable genes) and SSN data (i.e., network/edge biomarkers, top 100 variable edges) by package “ConsensusClusterPlus” in Bioconductor (http://www.bioconductor.org/). The log-rank *p* value of the survival curve was then used to evaluate the effect of the cancer subtyping. The numbers of subtypes for various cancers were referred to the following publications for BRCA (Reis-Filho and Pusztai, 2011), KIRC (Kim et al., 2002), LUAD (Hofree et al., 2013), ovarian serous cystadenocarcinoma (Hofree et al., 2013), and uterine corpus endometrial carcinoma (Hofree et al., 2013). As shown in Figure 6E, the accuracy (the *p* value) of subtype identification by our edge biomarkers (i.e., differential Δ*PCC*s) is superior to the traditional node biomarkers (i.e., differential genes), which demonstrates that the information of associations on an individual basis is useful and powerful in subtype identification of complex diseases.

### The functional driver genes from SSN

A mutation driver is the mutation that can confer growth advantage to the cell and be positively selected on cancer occurrence (Stratton et al., 2009). However, known DMGs are mutated in many cancer samples, and the existing driver mutations cannot explain all cancer onsets and typically have low coverage in cancer samples. Thus, in this work, we developed the concept of functional driver genes, whose dysfunction will benefit the formation or maintenance of the cancer occurrence with or without somatic mutations. The dysfunction can be measured by the changes in associations or network between the reference samples and the single sample, and thus the functional driver genes can be obtained by analyzing the SSN.

The high-degree genes in the SSN are important features for a cancer sample, and characterize their importance in the dysfunctional network or SSN of the single sample. Actually, they are also strongly related to the mutation driver genes in the individual sample (Figure 5E). Thus, the high-degree genes in the SSN can be considered as a measurement of the functional driver genes from the network viewpoint, and they may play an important functional role in the SSN for cancer occurrence and have the ability to drive the normal cell to a cancerous phenotype, similarly to mutation driver genes.

SSNs can be used to detect the functional drivers solely based on the expression data, even without the sequence information (i.e., without mutation information). We collected all of the top 5 highest degree genes, which are regarded as the potential functional driver genes of each single sample, for every sample in various cancers. The genes that appeared in at least 5 samples were chosen as the functional driver genes of this kind of cancer (Table S4), and were further validated as the potential disease genes by their mutation ratio in the cancer genomic data of TCGA with cBioPortal (Cerami et al., 2012). That is, we compared the ratio of the number of samples with mutations in these potential disease genes against the number of total samples (blue color, Figure S9), and the ratio of the number of samples with mutations in random genes (with the same number as the potential disease genes) against the number of total samples (orange color, Figure S9). The result shows that there is an obviously higher mutation ratio for these functional driver genes than random genes in cancer samples of TCGA. This result implies that the functional driver genes tend toward mutation or dysfunction in cancer samples.

### Experiments validated that SSNs identified functional driver genes contributing to drug resistance

Lung cancer is the leading cause of cancer-related deaths worldwide, with non-small cell lung cancer (NSCLC) being the predominant form of the disease (Siegel et al., 2014). The epidermal growth factor receptor (EGFR) signaling pathway, essential for normal epithelial cell proliferation, is frequently deregulated in lung cancer (Sordella et al., 2004). EGFR kinase inhibitors, including gefitinib and erlotinib, are clinically effective therapeutics for NSCLCs with EGFR kinase domain mutations (Ji et al., 2006; Mok et al., 2009). However, the clinical efficacy of gefitinib is limited by the development of acquired drug resistance. Using the human lung cancer cell line PC9 that harbors an EGFR kinase domain mutation and is sensitive to tyrosine kinase inhibitor (TKI) treatment, and the TKI-resistant cell line (PC9-DR) derived through long-term exposure to TKI, we performed microarray analyses of gene expression. The expression profiles of PC9 and PC9-DR were obtained separately, and then the two SSNs for PC9 and PC9-DR were both constructed based on the expression profiles and the reference dataset from GSE19804 of the Gene Expression Omnibus database (see Methods), and only the correlation-gained edges in the SSN were retained and used for the following analysis. Their differential network (Liu et al., 2012) was constructed by removing the common SSN edges of PC9-DR and PC9 from the SSN of PC9-DR, and 59 candidate genes as the potential functional driver genes were identified from the differential network based on the degree distribution (i.e., genes with degree > 10 in the differential network between the SSNs of PC9-DR and PC9) (Figure S10 and Table S5). Interestingly, most of these genes did not show significant expression changes between PC9-DR and PC9 and thus might not be identified by classical methods that are generally based on differential gene expression (Table S5). To test the power of our methods, we then performed individual gene knockdown of the top 15 candidates with small interfering RNA (siRNA) and analyzed their influence on cell growth and drug response (Figure S11).

Our data show that knockdown of these genes did not impact cell growth in both PC9 and PC9-DR cells (Figure S12). Interestingly, 73% of these genes (11 out of 15) were identified as important regulators of drug resistance since knockdown of any one of these 11 genes significantly conferred the PC9-DR cells with sensitivity to gefitinib (Figure 7B and C). Among them, five genes (BRCA1, GRB2, TGFBR1, CASP3, and CSNK2A1) showed the most significant effects when knocked down in PC9-DR cells (Figure 7B and C). Moreover, to compare with the analytical strategy that focuses on expression changes, we further knocked down the top 5 upregulated genes with short hairpin RNA (shRNA). However, none of these genes could overcome drug resistance when knocked down (Figure S13a). We also randomly choose 29 other genes to perform knockdown screening with siRNA, and PC9-DR cells showed no significant difference in drug response after their knockdown (Figure S13b). Taken together, our experimental data validate the effectiveness of the SSN method, and demonstrated the superiority of SSNs in the identification of genes important for drug resistance by considering individual networks. Our method is more powerful than traditional differential expression for identifying the functional driver genes.

**Figure 7.**
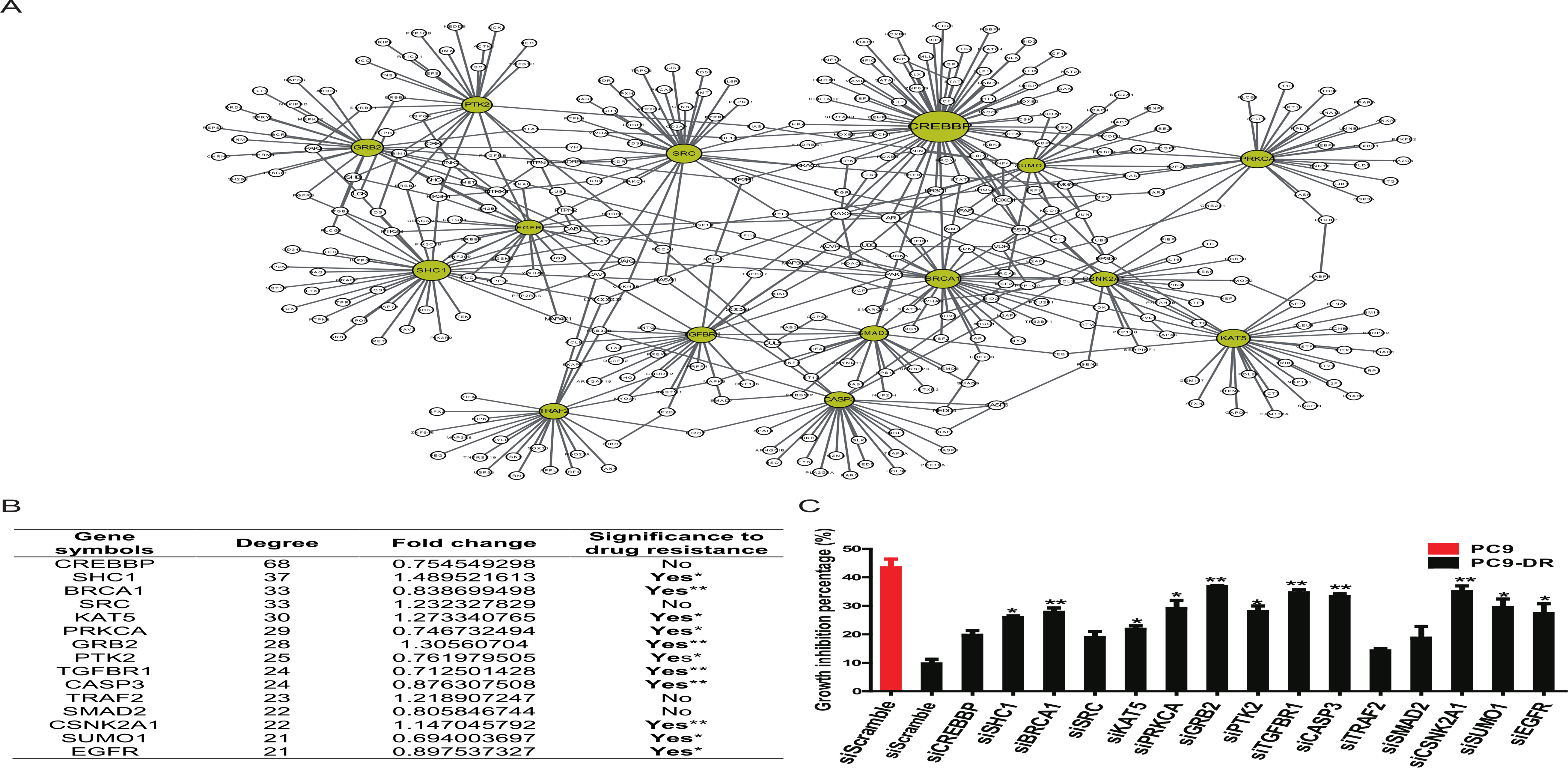
Experimental identification of functional driver genes contributing to drug resistance in the lung cancer cell line PC9-DR. (A) The subnetwork for the top 15 candidate genes. The node size indicates the degree of individual genes. The 15 candidate genes are highlighted as yellow, and other nodes are the first-order neighbors (genes) with the top 15 candidate genes in the differential network between PC9 and PC9-DR. The detailed differential network is provided in Figure S10. (B) The basic information for the top 15 candidate genes. Gene symbols: the official symbol for each gene; Degree: the degree of each gene in the differential network (or the number of neighbors for each gene); Fold change: the expression change of each gene from PC9 to PC9-DR; Significance to drug resistance: the significance of the results of drug resistance after gene knockdown. Clearly, none of the 15 genes exhibit significant differential expression (0.66<Fold change<1.5), and thus may not be identified by traditional statistical analyses, although most of them actually show significant effects on drug resistance (11 genes among 15 candidate genes) in knockdown experiments. (C) The growth inhibition percentage when either PC9 or PC9-DR cells with indicated gene knockdown were treated with 1 gefitinib for 72 h. PC9 siScramble was set as the positive control and PC9-DR siScramble was set as the negative control. Data are shown as means ±SEM. ^**^*p* < 0.01, \**p* < 0.05. Note that all edges shown in (A) are the upregulated edges (correlation-gained edges) from PC9 to PC9-DR.

### Theoretical relations between differential correlations and differential expression

We first define three types of edges in a network when there is a sample in addition to the reference samples. (a) The correlation-gained edge is the edge whose correlation or *PCC* in the absolute value is increased from the reference samples to the single sample, (b) the correlation-lost edge is the edge whose correlation or *PCC* in the absolute value is decreased from the reference network to the single sample, and (c) the correlation-invariant edge is the edge whose correlation or *PCC* in the absolute value exhibits little change from the reference network to the single sample.

We assume that there are two genes X and Y; the expressions of X and Y in the reference samples are *X_r_* and *Y_r_* with *r = 1,…, n;* the expressions of X and Y in the single sample is *X_S_* and *Y_S_;* the differential expression of X is 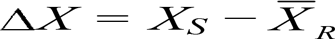, and the differential expression of Y is 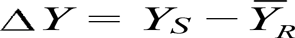, where 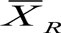 and 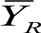 are average values of X and Y in the reference samples, respectively. Based on the analysis in Note S10, when the number of the reference samples, i.e., *n*, is sufficiently large, we can theoretically derive the following relations between differential correlation Δ*PCCn* and differential expression of X and Y:

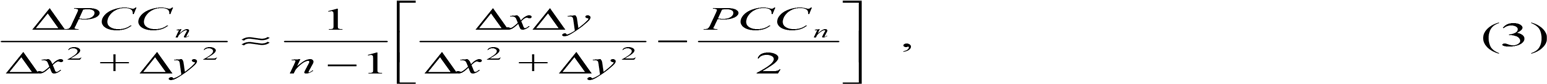

or

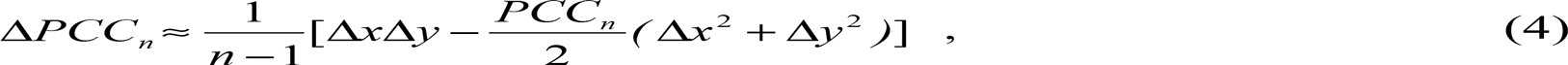

where Δ*x* and Δ*y* are Δ*X* and Δ*Y* normalized by the reference samples, defined as follows:

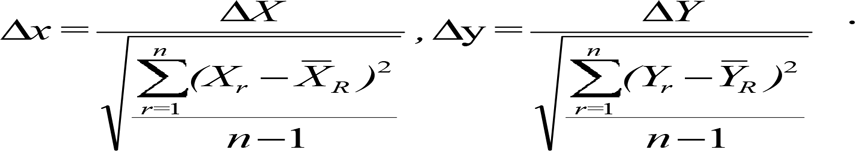

From the above equations (3) and (4), we can obtain Tables S6 and S7, which describe the various cases between Δ*PCC* and differential expression of X and Y, and a graphical explanation is also given in Figure S13. The details of the different gene expression levels for a single sample affecting the correlation are given in Supplemental Information Notes S10 and S11 with Tables S6 and S7.

## DISCUSSION

The Δ*PCC* is the interferential degree of correlation of a single sample perturbing the reference samples, and it depicts the changed degree of correlation by adding the single sample to the reference samples. Thus, it describes the difference in associations from a network viewpoint. If the single sample can affect the correlation of two genes in the reference samples with a significant change, the regulation of two genes in the single sample is considered to be inconsistent with the regulation in the reference samples. This inconsistency of regulation may be due to the differential gene expression in either or both of the two genes, or caused by a functional alteration, e.g., a mutation that cannot be identified by traditional testing of differential expression of the genes. Therefore, Δ*PCC* testing is a more sensitive method than the traditional differential expression testing, and can identify the potential disease genes that even display no differential expression from normal/control samples. In such a sense, SSN is complementary to the traditional methods from the network perspective.

A biological function is generally facilitated not by individual molecules but by their regulations or molecular networks, which dynamically change with time and conditions. Thus, identifying the condition-specific network or SSN is crucial to elucidate molecular mechanisms of complex biological processes at a system level. However, although expression data or sequencing data provide information about the profiles of molecules on a single-sample basis, there is no effective method to construct a molecular network on a single-sample basis. In this work we proposed a statistical method to construct the SSN for a single sample, which opens a new way for both characterizing personalized features and analyzing biological systems at a network level. The analyses of TCGA data not only validated the effectiveness of our method but also demonstrated that SSNs can characterize the network patterns on a single-sample basis. We also reported new discoveries for regulatory patterns, personalized networks, and edge biomarkers in several cancer types.

Although a group of reference samples is required in our method, it is generally available even in clinical practice and also there is no strict condition on the reference samples. Theoretically, the reference samples can be composed of any type of sample, but choosing those reference samples with distinct expression profiles from the test samples certainly increases the discriminatory power of its SSN. Actually, to check the robustness of the results against the different choices of the reference samples, we tested breast cancer data from TCGA. There are 99 normal samples from TCGA, and we randomly chose 17 normal samples from these as a group of reference data. With these 17 randomly chosen reference samples, we could then construct the SSN for each cancer sample by our method and compared the new SSNs with the old SSNs. We repeated the process 100 times, and obtained the average recurrence ratio of the edges in SSN from different reference samples. The comparison results show that the average recurrence ratio of the edges in the SSN from the different reference samples is as high as 81.01%, which indicates that the method is stable and robust with respect to the choice of the reference (or normal) samples. Another test was also performed to check the robustness of the SSN from different reference sample sizes based on a breast cancer dataset. We randomly chose 15, 20, 30, and 50 normal samples from 99 normal samples as reference samples, and calculated the SSN for all tumor samples. The new SSNs were then compared with the old SSNs from 17 control samples, and this test was repeated 100 times. The average recurrence rate of edges in the new SSNs relative to the old SSNs from 17 control samples is 80.34, 82.38, 83.8, and 85.17% for reference sample sizes 15, 20, 30, and 50, respectively. These results indicate that the method is robust and stable for the different reference sample sizes. The high-degree genes in the SSN are important features for a cancer sample, and are strongly related to the DMGs in the individual sample, which may be beneficial to personalized diagnosis or individualized treatment. We show that the high-degree genes of an SSN can be used to predict the DMGs for this sample, and the accuracy of the prediction for the DMGs increases with the degree of the gene in SSNs. This property implies that SSNs can be used to detect the functional drivers solely based on the expression data even without the sequence information.

Generally, there are three types of perturbation for a hub in an SSN, i.e., a hub in an SSN could be the result of (a) a perturbation in the hub gene itself, (b) a perturbation of its interaction partners, or (c) a combination of both, although there are few such hub genes for the type “(a)” among TCGA samples. One possible reason is that a gene is generally connected with other upstream and downstream genes in the form of a feedback network, which implies few such cases in biological systems. On the other hand, types “(b)” and “(c)” are widely observed in TCGA samples. Even for one hub gene in the same disease, there are both types “(b)” and “(c)”. For example, TP53 has no differential expression in samples A0HO and A0DP in breast cancer (Figure 3A), i.e., TP53 is the type “(b)” in these two samples, but in samples A0B0 and A25A, TP53 is overexpressed relative to the control samples (Figure 3A), i.e., TP53 is the type “(c)” in these two samples. In particular, there are 37531 hub genes with high degrees (i.e., at least 10 neighbors) in SSNs of 761 breast cancer samples, where no sample is of type “(a)”, 93 of 37531 hub genes belong to type “(b)”, and most of the hub genes are of type “(c)”.

It should be noted that an SSN in this work is not a real molecular network for each sample but a perturbation network for a single sample against the reference network. It reflects the variation between normal and disease samples in terms of interactions, regulations, or a network, similarly to differential expression of a gene, which is not the real gene expression level for each sample but the variation of the gene expression between normal and disease samples. In contrast to a molecular network inferred by traditional methods, which is actually an aggregated network for multiple samples, our method can construct an SSN on a single-sample basis and thus can be directly applied to the data analysis of single samples, in particular with potential applications to precision medicine and personalized medicine.

This method is based on the *PCC*, and thus the correlation network is the desired choice to construct the SSN. However, humans have more than 20,000 genes, which implies that the full-correlation network for humans has more than 200,000,000 edges. Hence, to construct such a network is computationally intense; in addition, the correlation network includes indirect associations, which are false-positive connections in a molecular network. In this work, we adopted the background or reference network to reduce the false-positive connections and also significantly alleviate the computational and storage requirements.

Some recent studies (Bhardwaj and Lu, 2005; De Bodt et al., 2009; Liu et al., 2012; Zhang et al., 2013) were developed to decompose the aggregated associations of a group of samples into those of individual samples. This method has some similarities with ours but is notably different. In particular, the previous method (Bhardwaj and Lu, 2005; De Bodt et al., 2009; Liu et al., 2012; Zhang et al., 2013) approximately decomposes the association or *PCC* into a group of networks corresponding to individual samples. There are three major differences between those two methods. First, an SSN in this work is actually a perturbed network of an individual sample from a group of reference samples, which characterizes the individual sample at the network level in an accurate manner. In contrast, the previous method (Bhardwaj and Lu, 2005; De Bodt et al., 2009; Liu et al., 2012; Zhang et al., 2013) uses an approximation scheme to decompose an association (or aggregated) network of a group of samples approximately into individual networks corresponding to individual samples in a linear manner. Second, our method evaluates a new single sample based on a group of reference samples, whereas the previous method (Bhardwaj and Lu, 2005; De Bodt et al., 2009; Liu et al., 2012; Zhang et al., 2013) evaluates a single sample in the group. Third, an SSN is constructed using a statistic based on a new type of distribution, the volcano distribution, which can be proven mathematically and ensured by statistical theory. However, the previous method (Bhardwaj and Lu, 2005; De Bodt et al., 2009; Liu et al., 2012; Zhang et al., 2013) uses no such statistic, and furthermore the correlation network constructed by the previous method (Bhardwaj and Lu, 2005; De Bodt et al., 2009; Liu et al., 2012; Zhang et al., 2013) may be improper both theoretically and numerically, i.e., the correlation in each individual network can be more than 1 or less than −1 due to its heuristic scheme (see Eqn. (31) in the references (Bhardwaj and Lu, 2005; De Bodt et al., 2009; Liu et al., 2012; Zhang et al., 2013)).

Biological experiments on drug resistance validated not only the effectiveness of our method for constructing SSNs by single samples, but also one advantage of our method, i.e., identifying those non-differentially expressed disease genes or factors as functional drivers, which are generally missed by traditional methods. As shown in Figure 7, although there are no differentially expressed genes between PC9-DR and PC9 in the top 15 candidate functional driver genes, the expression of some genes was actually reduced from PC9 to PC9-DR (Figure 7B) (not significant in terms of fold change). After knocking down these downregulated genes, i.e., BRCA1, TGFBR1, and CASP3, the drug resistance of PC9-DR showed very significant changes (Figure 7B and C). This result cannot be explained by traditional analysis based on differential expression, but in our work, we show that the correlations of these genes with their neighbors were increased from PC9 to PC9-DR, i.e., the regulation between these genes and their neighbors increased with the drug resistance, even though their gene expression decreased. Thus, deeply knocking those genes down reduced the regulation with their neighbors and therefore changed the drug resistance.

In this work, we constructed an association network by correlation, which includes both the effect of direct and indirect regulation between two genes. Actually, instead of the correlation or *PCC*, we can similarly use the partial correlation or partial mutual information to construct a direct association network (Zhang et al., 2015), which we will address in future studies.

## METHODS

The procedure to construct an SSN is shown in Figure 1, and the theoretical derivation and explanation are given in Results and Supplemental Information Notes S2-S4. In this paper, SSN also implies an individual-specific network.

### Topological distance between genes in the SSN and SMGs in the same sample

Let the genes in the SSN be a set *S*, and the SMGs in this single sample be a set *D*. The topological distance is based on the shortest distance between these two sets *S* and *D* in the background network. Specifically, for a single sample, the average shortest distance was calculated by averaging the shortest distances between each gene on the SSN and each SMG of this sample based on the connection of the background network, where the “igraph” extension module (http://igraph.sourceforge.net/) of the Python programming language was used to obtain the shortest distance between two genes in the network. If two genes cannot be linked based on the background network, then the shortest distance between the two genes is assigned 100, which is a sufficiently large value for the shortest distance in the network. Then, the same number of genes of the SSN were randomly chosen from a background network for this single sample, and the average shortest distance between these randomly chosen genes and the SMGs of this sample were again computed by “igraph”, and the average shortest distance for the random genes was compared with the average shortest distance for the genes in the SSN. This permutation was repeated 100 times. The proportion, in which the average shortest distance for random genes is less than the average shortest distance for the genes in the SSN, is defined as the topological distance between the SSN and the SMGs for this single sample. If the proportion is less than 0.05, we consider the topological distance to be significant for this sample. Otherwise, the topological distance is not significant.

### Functional distance between the SSN and the SMGs in the same sample

For a single sample, the pathway or functional enrichment of the genes in the SSN based on the KEGG or GO pathway was calculated by the hypergeometric test (Rivals et al., 2007) as follows:

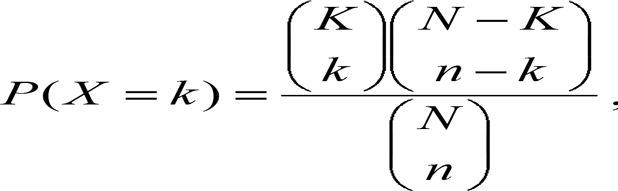

where *N* is the number of genes on the whole background network, *K* is the number of genes sharing a pathway or a GO term on the background network, *n* is the number of genes in the SSN of a single sample, and *k* is the number of genes sharing the same pathway or GO term in the SSN, *P*(*X* = *k*) is the probability for observing exactly *k* shared genes in the hypergeometric distribution. The *p* value of the hypergeometric distribution is calculated by the cumulative probability *P*(*X* ≥ *k*) (http://en.wikipedia.org/wiki/Hypergeometric_distribution). If the *p* value of the enrichment for a pathway or a GO term is less than 0.05, then we regard that this pathway or GO function is significantly enriched in the SSN of this single sample. Otherwise, we regard that the pathway or GO function is not enriched in the SSN. Subsequently, the functional association of the SSN and somatic mutations in the same sample can be defined as the number of shared pathways in KEGG or functions of the GO terms between the enriched pathways or GO terms of the SSN and each SMG. The genes with the same number as that of the SSN were then randomly chosen from this single sample, and the enriched pathways or GO terms were again identified by the hypergeometric test. The functional association between these randomly chosen genes and the SMGs of this sample were separately obtained based on KEGG and GO, and the random functional association was compared with the actual functional association between the SSN and somatic mutations of this sample. This permutation was repeated 1000 times. The proportion, in which the random functional association is less than the actual functional association, is defined as the functional distance between the SSN and SMGs for this single sample. If the proportion is less than 0.05, we consider the functional distance to be significant for this sample; otherwise, the functional distance is not significant for this sample.

### Classification of phenotypes and subtypes of cancer

A five-fold cross-validation was conducted for the classification of phenotypes by the “ksvm” package in Bioconductor for the R language to implement the function of SVM, and the ROC curve is drawn by the “ROCR” package in Bioconductor for the R language. The hierarchical clustering was also used to classify the phenotype in the R language.

For the subtype of cancer, the top 100 nodes (differentially expressed genes) or edges (Δ*PCC*s) with standard deviations were chosen for the consensus clustering, and the “ConsensusClusterPlus” package in Bioconductor for the R language was used to perform the consensus clustering. The “survival” package in Bioconductor was used to calculate the log-rank value of the survival curve.

### Cell culture and siRNA transfection

The NSCLC cell line PC9 expressing the EGFR exon 19 deletion mutation was purchased from ATCC (Manassas, VA, USA) and was grown in RPMI 1640 medium supplemented with 10% fetal bovine serum. The gefitinib-resistant cell line derived from PC9 (PC9-DR) was established by treating PC9 cells with gefitinib continuously for 3 months. siRNA transfection was performed using RNAimax (Life Technologies, Carlsbad, CA, USA) following the manufacturer’s protocol. Two siRNAs were used for each gene.

### Constructing SSNs for PC9 and PC9-DR

The expression profiles of PC9 and PC9-DR were detected by a Human Genome U219 Array (Affymetrix, Santa Clara, CA, USA), and 2 repeats for both PC9 and PC9-DR were performed in this study. Sixty normal samples were chosen from the GSE19804 dataset (http://www.ncbi.nlm.nih.gov/geo/query/acc.cgi?acc=GSE19804) as the reference samples for constructing the reference network. Two PC9 SSNs and two PC9-DR SSNs were then separately constructed based on the reference network and the expression profiles of PC9 and PC9-DR. We only chose the correlation-gained edges from these SSNs for the functional validation experiments (due to gene knockdown experiments). After filtering the correlation-lost edges, the overlapped network of two PC9 SSNs was taken as the SSN of PC9, and the overlapped network of two PC9-DR SSNs was taken as the SSN of PC9-DR. Subsequently, the differential network (Liu et al., 2012) between the SSNs of PC9 and PC9-DR was constructed by subtracting the SSN of PC9 from that of PC9-DR (i.e., removing the common SSN edges of PC9 and PC9-DR from the SSN of PC9-DR). The high-degree (> 10) genes in the differential network were then selected as potential genes for drug resistance. Clearly, all edges in this differential network are the upregulated edges (correlation-gained edges) from PC9 to PC9-DR.

### Drug treatment and cell growth assay

Cells were plated in triplicate at a density of 3000 cells/well in 96-well plates. The cells were then treated with 1 µM gefitinib for 72 h and the cell growth assay was performed as previously described (Li et al., 2015).

### shRNA experiments

The shRNAs against the top 5 upregulated genes (FAM171A1, COL13A1, vimentin, BMP5, and CYB5R2) were subcloned into the pLKO.1 vector (Addgene, Cambridge, MA, USA). shRNA against luciferase gene was used as control. shRNAs were packaged in lentiviral particles by cotransfecting with packaging plasmids into 293T cells and the filtered cell culture supernatant was further used to infect PC9-DR cells as previously reported(Gao et al., 2014). 293T cells were cultured in DMEM with 8% FBS. The shRNA sequences used are listed in Table S8.

### Reverse transcription PCR and quantitative real-time PCR (qPCR)

RNA was extracted using Trizol reagent (Invitrogen, Carlsbad, CA, USA) and phenol/chloroform methods and then reverse-transcribed into first-strand complementary DNA with a RevertAid First Strand cDNA Synthesis Kit (Fermentas, Waltham, MA, USA). Gene overexpression and knockdown efficiency were detected by qPCR with gene-specific primers using a 7,500 Fast Real-Time PCR System (Applied Biosystems, Foster City, CA, USA) and SYBR Green Master PCR Mix (Invitrogen). Glyceraldehyde 3-phosphate dehydrogenase (human) served as an internal control. The primers used for PCR are listed in Table S8.

### Statistical Analysis

Statistical analysis was performed using a two-tailed Student’s *t* test.

## Authors’ Contribution

L. Chen and K. Aihara planned this study; L. Chen, H. Ji, X. Liu, and Y. Wang designed the experiments; X. Liu performed the dry experiment; Y. Wang performed the wet experiment; X. Liu, Y. Wang, and L. Chen wrote the manuscript; and L. Chen, H. Ji and K. Aihara revised the manuscript.

## Acknowledgments

This work was supported by the Strategic Priority Research Program of the Chinese Academy of Sciences (No. XDB13040700), the National Program on Key Basic Research Projects (No. 2014CB910504, 2012CB910800), the National Natural Science Foundation of China (NSFC) (Nos. 61134013, 91439103, 61403363, 81430066, 81325015, 31370747, 81101583, 81372509) and Science and Technology Commission of Shanghai Municipality (12JC1409800). This work was supported by the Knowledge Innovation Program of SIBS of CAS (No. 2013KIP218) and China Postdoctoral Science Foundation (Nos. 2014T70441, 2013M541565). This work was also supported by JSPS KAKENHI Grant Number 15H05707 and JST’s “Super Highway”, the accelerated research program to bridge university IPs and practical use. The authors also appreciate the valuable suggestions of Dr. Dangsheng Li on both the experiments and the manuscript.

